# A novel antioxidant N-acetylcysteine Amide Alleviates Cyclophosphamide-induced Endothelial Damage

**DOI:** 10.1101/2025.04.07.647539

**Authors:** Rui He, Wenyi Zheng, Terra Slof, Eva Wärdell, Agneta Månsson-Broberg, Svante Norgren, Ying Zhao, Moustapha Hassan

## Abstract

The alkylating agent cyclophosphamide (Cy) is one of the important corner stones in cancer treatment. Cy is used also as a part of conditioning regimens prior to hematopoietic cell transplantation and as a prophylactic treatment post transplantation in graft-versus-host disease. Existing evidence showed that high doses of Cy are associated with a number of side effects, including damage on arterial endothelium, which might contribute to late cardiovascular disorders. Oxidative stress has been characterized in such pathogenesis and is an exploitable target for treatment. Herein, the study aimed to investigate the protective role of the novel antioxidant N-acetylcysteine amide (NACA) in Cy-induced endothelial injury and explore the underlying mechanism. Our *in vivo* results showed that NACA partially reduced the endothelial injury and recovered the integrity of arterial endothelium in the mice treated with Cy. In addition, we found that NACA decreased the cytotoxicity of Cy on endothelial cells through alleviating caspase-dependent apoptosis, DNA damage and oxidative stress. Meanwhile, NACA pre-treatment rebalanced endothelial nitric oxide synthase (eNOS) and arginase I and preserved the angiogenic capability of endothelial cells which was compromised by Cy through blockage of Notch signaling pathway. Interestingly, in comparison to N-acetylcysteine (NAC), its amide derivative NACA showed superior ability to alleviate Cy-induced endothelial damage. In conclusion, the current study proved the robust endothelial protective potential of NACA, facilitating clinical use of the novel antioxidant.

## Introduction

The alkylating agent cyclophosphamide (Cy) is a potent drug with a broad range of indications for cancer, such as lymphoma, breast cancer, ovarian cancer, and bone and soft tissue sarcomas[1-4], by causing irreversible DNA double strands breaks and cell apoptosis. Apart from the anti-neoplastic effect, Cy has been widely used in the conditioning regimens prior to hematopoietic cell transplantation (HCT) and as a prophylactic treatment post transplantation in graft-versus-host disease (GvHD) through its immunosuppressive property. However, the doses of Cy used in HCT-related applications are extremely high, with a conventional dose of >120 mg/kg administered over 2–4 days as conditioning[5], and a dose of 50 mg/kg/d given on days 3 and 4 after transplantation as prophylaxis[6]. Such high-dose Cy treatment is associated with serious adverse effects, including alopecia, gonadal toxicity, hemorrhagic cystitis, carcinogenesis, and cardiac dysfunction. It has been reported that myocarditis, pericardial effusion, pericarditis and heart failure occurred in 25% of cases at a dose of ≥ 1.55g/m^2^/d[7], which is corresponding to a dose around 46 mg/kg/d.

The manifestations of Cy-induced cardiotoxicity are heterogeneous and range from innoxious to lethal[8], and the clinical presentations include myocardial necrosis, myocarditis, cardiomyopathy, etc[9]. Since the myocardium mainly consists of cardiomyocytes, cardio fibroblasts, and endothelial cells[10], the cardiotoxicity caused by Cy may take place in all three cell types. However, tremendous research efforts have focused on the damage of cardiomyocytes[11, 12], while endothelium with the errand to maintain homeostasis and vascular tone has been poorly studied. Given that endothelial cells are more susceptible to Cy-induced damage than other cells [13] due to its higher turnover rate[14], and that early endothelial dysfunction is predisposes to atherosclerosis which can later on result in fatal cardiovascular events[15], it is of great importance to explore strategies to preserve the integrity of cardio and arterial endothelium.

Limited evidence showed that Cy-mediated endothelial dysfunction involves different molecular mechanisms including reduction in progenitor endothelial cells, endothelial nitric oxide synthase (eNOS) uncoupling, compromised nitric oxide (NO) bioavailability, increased oxidative and nitrative stress, overproduction of endothelin-1 (ET-1), endothelium inflammation, thrombosis, etc[16-19]. Among them, elevated oxidative stress was found to be the major contributor and associated with other mechanisms. Therefore, approaches to counteract oxidative stress was assumed to attenuate Cy-induced endothelial injury.

N-acetylcysteine (NAC) is a classic antioxidant and clinically prescribed as an antidote for paracetamol overdose and as a mucolytic agent[20, 21] due to its ability to replenish glutathione (GSH) synthesis as well as scavenge reactive oxygen species (ROS). However, its efficacy in combating intracellular oxidative stress is limited by poor permeability across cellular membrane. N-acetylcysteine amide (NACA) was subsequently synthesized with significantly improved lipophilicity and membrane permeability. Preclinical studies[22, 23] have indicated its robust antioxidative ability and tolerability *in vitro* and *in vivo*. In our recent pharmacokinetic study, NACA was proved to be superior to NAC in terms of oral bioavailability (67% vs 15%) and capacity to replenish GSH[24]. Taken together, we assume that NACA might attenuate Cy-induced endothelial damage with higher efficacy than NAC.

The current study sought to investigate the potential role of NACA in maintaining the integrity of vessel endothelium after exposure to Cy, especially in comparison to NAC. To reveal the underlying mechanisms of action, cellular morphology, cell death pathway, DNA damage, redox hemostasis, and tube formation ability were evaluated.

## Materials and methods

### Reagents

NACA was provided by Dr Glenn Goldstein, Sentient Life Sciences Inc, NY, New York, USA, while NAC (A7250-25G) and cyclophosphamide monohydrate (C0768-25G) were bought from Sigma-Aldrich (St. Louis, Missouri, USA). NACA and NAC were dissolved in normal saline or cell culture medium and naturalized with NaOH freshly before use. Mafosfamide (MAF, D-17272) from Niomech-IIT GmbH (Bielefeld, Germany) was prepared on ice at a concentration of 5 mM for further dilution.

### Animals

Animal experiments were approved by the Stockholm Southern Ethical Committee (ethical permit No. ID 11257-2020). NACA or NAC was orally administered to Babl/c mice at a dose of 250 mg/kg, twice a day (from day -7 to day -1 and from day 1 to day 7). Cy was intraperitoneally injected into the mice at a single dose of 400 mg/kg on day 0 (Figure 1A). Blood samples were collected and the cardiac troponin I (cTNI) levels in serum were measured using a commercial ELISA kit (Life diagnostics, CTNI-1-HS). Aortas were fixed and the paraffin sections were stained with H&E, and the histopathological structures were imaged using a bright-field microscope (Nikon, NIS-elements, F 3.0) at 400× magnification.

**Figure 1.**
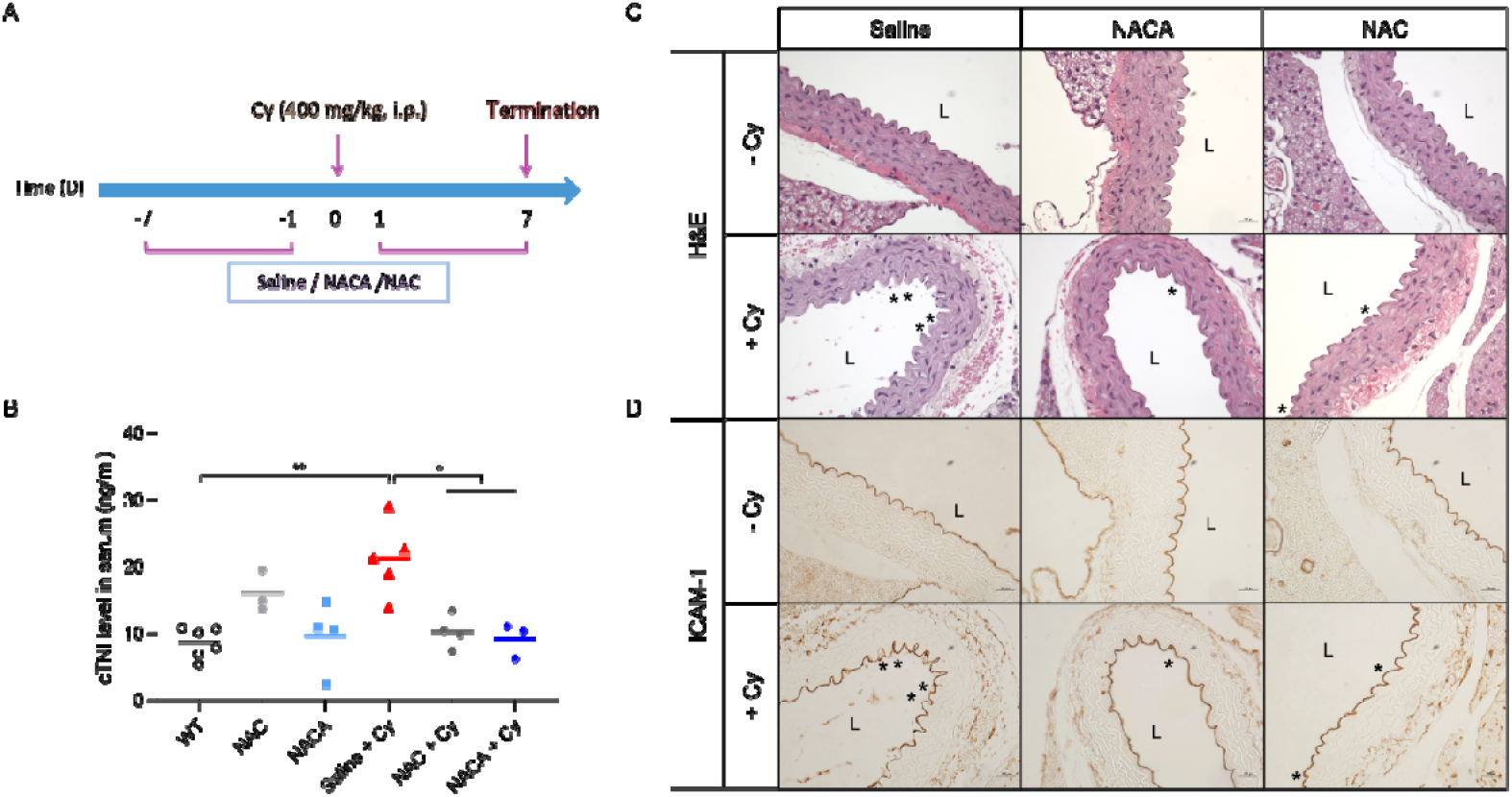
NACA mitigated cardiac and endothelial damage caused by Cy. (A) Schedule of treatments on Balb/c mice. NACA or NAC was orally administered to Balb/c mice at a dose of 250 mg/kg, twice daily (day -7 to day -1, day 1 to day 7). Cy was intraperitoneally injected into the mice at a single dose of 400 mg/kg on day 0. (B) Expression levels of cTNI in the mouse serum. Mice were sacrificed after the last dose of NACA or NAC. cTNI levels in serum were measured using a commercial ELISA kit. Results were presented as individual data with mean value of each group. *: p < 0.05; **: p < 0.005. (C) Representative H&E staining on the murine aortic tissue. L: lumen; *: suspected damaged site. (D) Representative immunohistochemical staining with ICAM-1 antibody. *: where ICAM-1 protein was highly expressed.

### Immunohistochemical staining

Paraffin-embedded aorta sections were rehydrated, and antigens were retrieved using a citrate buffer (Sigma-Aldrich, C9999) as the unmasking solution. To eliminate endogenous peroxidase activity, tissues were placed in 3% H_2_O_2_ for 20 min at 37 °C. After blocking, sections were incubated with Recombinant Anti-ICAM1 antibody (1:500, Abcam, ab179707) overnight at 4 °C, followed by HRP-labelled secondary antibody (1:200, Abcam, ab6721) incubation at RT for 30 min. DAB substrate kit (Abcam, ab64238) was applied to visualize the antigen expression.

### Cell culture

Human aortic endothelial cells (HAECs) and human umbilical endothelial cells (HUVECs) were purchased from Promocell GmbH (Heidelberg, Germany). All cells were maintained at 37 °C with 5% CO_2_ and cultured in endothelial cell media MV2 (Promocell, C22022) supplemented with penicillin-streptomycin (Sigma-Aldrich, P4333). Cells from passages 2-10 were used for experiments, except for tube formation assay, in which cells with passages 3-4 were utilized.

### Cell proliferation assay

Endothelial cells were seeded in a 96-well plate (6×10^3^ cells/well) and a 12-well plate (72×10^3^ cells/well) and left to attach overnight. After pre-treatment with NACA or NAC (1mM or 5 mM) for 6 hr, the cells were rinsed with Hank’s Balanced Salt Solution (HBSS, Gibco) and further treated with MAF solution for 20 hr. Cell viability was assessed by WST-1 kit (Sigma-Aldrich, A7250) and crystal violet staining.

### Flow cytometry

HAECs and HUVECs were seeded in 6-well plates (2×10^5^ cells/well) and treated as mentioned above. Cells were harvested by gently trypsinization. Apoptosis was analyzed using PE-Annexin V apoptosis detection kit I (BD, 559763) according to manufacturer’s instructions, and data was obtained by a flow cytometer (FACS Array, BD). For DNA damage assay, cells were fix with 80% methanol for 5 min at room temperature (RT) and rinsed with ice-cold Dulbecco’s Phosphate Buffered Saline (DPBS, Gibco). Then the cells were permeabilized with 0.1% Triton-X (10 min, RT) and blocked in 3% BSA for 1 hr at RT. Thereafter, cells were incubated with recombinant anti-γH2A.X (phospho S139) antibody (1:200, Abcam, ab81299) overnight at 4°C, followed by 1-hour incubation with secondary FITC-labelled antibody (1:1000, Abcam, ab7086). Cells were then analyzed on a flow cytometer (CellStream™, Merck).

### Determination of reactive oxygen species

Cellular ROS level was evaluated by a fluorescence-based kit (Abcam, ab13851). Briefly, cells were seeded in a 96-well plate (8×10^3^ cells/well) and pre-incubated with NACA or NAC for 4 hr. HAECs or HUVECs were stained with DCF-DA for 45 min, and then treated with 85μM or 100 μM MAF for 3 hr, respectively. The fluorescence intensity was quantified using a microplate reader. Phenol red-free cell culture medium (PromoCell, C-22226) was used to avoid interference.

### Enzyme activities

HAECs and HUVECs were seeded in petri dishes (1×10^6^ cells/dish) and treated as mentioned above. After harvesting, cells were lysed by sonication. Cellular catalase and total superoxide dismutase (SOD) activity were measured using commercial kits (Invitrogen, EIACATC and EIASODC) according to methods described in the manufacturer’s instructions. Protein concentration of each sample was detected using BCA protein assay (ThermoFisher, 23227) and the enzyme activities were normalized basing on the total protein concentration.

### Quantification of reduced glutathione

After treatment, cells were lysed with mammalian cell lysis buffer (Abcam, ab179835) and protein concentration was evaluated. To exclude the enzymes that can interfere with the assay, protein was precipitated using a deproteinizing sample preparation kit (Abcam, ab204708). Cellular reduced GSH level was determined using a commercial assay (Abcam, ab205811) following manufacturer’s instructions. Fluorescence was monitored at Ex/Em = 490/520 nm with a microplate reader.

### Tube formation assay

Growth factor reduced basement membrane extract (BME) matrix (PromoCell, PK-CA577-K518-5) was thawed at 4 °C, pipetted into a pre-cooled 96-well plate, and incubated at 37 °C for 1 h. After BME gels polymeration, NACA or NAC pre-treated cells or control cells were suspended in MV2 medium with MAF and were seeded onto the gels. After 24 hr, the tubular structures were randomly imaged by a phase-contrast microscope (Olympus, CellSens software 1.16) at 40× magnification.

### Western blot

Cells were lysed in RIPA buffer (CST, 9806) supplemented with protease and phosphatase inhibitor (CST, 5872). Protein concentration was measured using BCA protein assay. Samples were denatured at 95°C for 5 min and equal amounts of protein were loaded on 4-15% Mini-protean^®^ TGX™ precast gels (Bio-Rad, 456-1085). Electrophoresis and transfer were performed by Bio-Rad systems. Membranes were blocked at RT for 1 hr in 5% (w/v) non-fat milk (Bio-Rad, 1706404XTU) dissolved in 1X Tris-buffered saline containing 0.1% (v/v) Tween 20 (TBST) and were incubated with primary antibodies at 4°C overnight. After staining with fluorescent secondary antibodies (1:5,000, Li-cor, IRDye ^®^ 680RD/800CW donkey anti-mouse/rabbit IgG), protein bands were visualized using Odyssey CLx (Li-cor) system. The following primary antibodies were employed: Actin (1:5,000, Sigma-Aldrich, A5441,), Caspase-3 (1:500, CST, 9662S,), Caspase-9 (1:500, CST, 9502S,), eNOS (1:300, Abcam, ab76198,), Arginase I (1:500, Invitrogen, PA5-85267,), Arginase II (1:1,000, Invitrogen, PA5-78820,), Jagged-1 (1:300, CST, 2620S), HES-1 (1:500, ThermoFisher, PA5-28802), Notch-1 (1:500, CST, 3608S).

### Quantification and statistics

The tube formation assay was analyzed and interpreted using ImageJ software (version 2.0), while the results from western blot were summarized by Image Studio Lite software (version 5.2.5). All data are presented as means ± SD (standard deviation) unless otherwise described. The statistical analyses between two groups were performed using Mann-Whitney t test, and P values < 0.05 were considered statistically significant.

## Results

### *In vivo* protective effect of N-acetylcysteine amide

Early endothelial dysfunction is associated with late cardiac injury, such as atherosclerosis, recurrent ischemia, and heart failure[15, 25]. As circulating cTNI is a valid biomarker for myocardial infraction and heart failure that could be detected at early stage, it may play a role in the recognition of endothelial dysfunction[26]. Thus, we examined the level of cTNI in the mouse serum to assess the ability of NACA in reducing Cy-induced endothelial damage. As shown in Figure 1B, Cy treatment greatly increased the level of cTNI in the serum compared to the WT group (21.3 ± 5.5 ng/ml vs 8.7 ± 2.3 ng/ml). Moreover, pretreatment with NACA and NAC significantly decreased the level of cTNI, to a concentration of 9.2 ± 2.6 ng/ml and 10.3 ± 2.5 ng/ml, respectively. However, there was no statistically significant difference between NACA and NAC pretreated groups.

H&E staining was performed to visualize possible damage in aortic endothelium. For the mice without Cy treatment, endothelial nuclei were apparently present in the inner layer of the vessels, and the tunica media consisted of several layers of elastic lamina was supported by arranged smooth muscle cells and linked to tight junctions with tunica intima. Administration of 400 mg/kg Cy caused conspicuous detachment of the endothelial nuclei along with an irregular arrangement of the matrix fiber and smooth muscle cells in the middle elastic lamina. Both NACA and NAC treatments showed the ability to restrain the impairment, although with different degrees (Figure 1C).

We further investigated the expression level of intercellular adhesion molecule-1 (ICAM-1) which marks endothelial dysfunction and determines inflammatory responses of endothelium[27, 28]. The results revealed that prophylactic treatment with NACA decreased the aortic endothelial ICAM-1 protein levels which was supposed to be upregulated by high-dose Cy. NAC-pretreated mice had less decrease in the level of ICAM-1expression (Figure 1D).

### Cell morphology and viability

A range of *in vitro* studies were conducted on human endothelial cells to better understand the courses by which Cy did harm to the vascular endothelium and the pathways through which NACA exerted its protective ability. As Cy is a prodrug and is metabolized by cytochrome P450 enzymes (CYPs) to the main active and cytotoxic metabolite, 4-hydroxycyclophosphamide (4-OH-Cy), MAF was used as the parent compound to produce 4-OH-Cy through automatic hydrolysis in aqueous phase due to the lack of major CYPs in human endothelial cells.

To figure out the appropriate working concentration of 4-OH-Cy, its cytotoxicity was determined on human endothelial cells. After treatment for 20 hr, 4-OH-Cy had an IC50 of 32 μM on HAECs and 36 μM on HUVECs, respectively (Figure S1A). At the concentrations around IC50, the toxicity of 4-OH-Cy shifted drastically despite of subtle changes in the concentration. To detect the protective effect of antioxidants in a reproducible manner, the acquired IC50s were thereafter applied for further investigation. We also examined the cytotoxicity of NACA and NAC on HAECs and HUVECs. Both substances showed no toxic effect on the cells at a concentration of 1 mM and 5 mM (Figure S1B and 1C). Therefore, we selected both antioxidants at 1 mM and/or 5 mM in our subsequent studies for comparison.

The morphological alterations of HAECs and HUVECs were first investigated. As expected, untreated cells showed flat and stretched shapes, clear cell boundaries and well-adherent stream-like growth, while 4-OH-Cy induced rounding and shrinking of endothelial cells. Notably, pre-treatment with NACA and NAC partially preserved the cell morphology, with 5 mM of NACA having the best protective effect (Figure 2A and 2B). Next, we performed violet staining and cell viability assay to quantify cell mass. It is shown that both NACA and NAC could prevent endothelial cells from the toxicity caused by 4-OH-Cy in a concentration-dependent manner. Most importantly, NACA showed a significant improvement compared to NAC at the same concentration, and nearly all cells were retrieved with 5 mM of NACA (Figure 2C-F).

**Figure 2.**
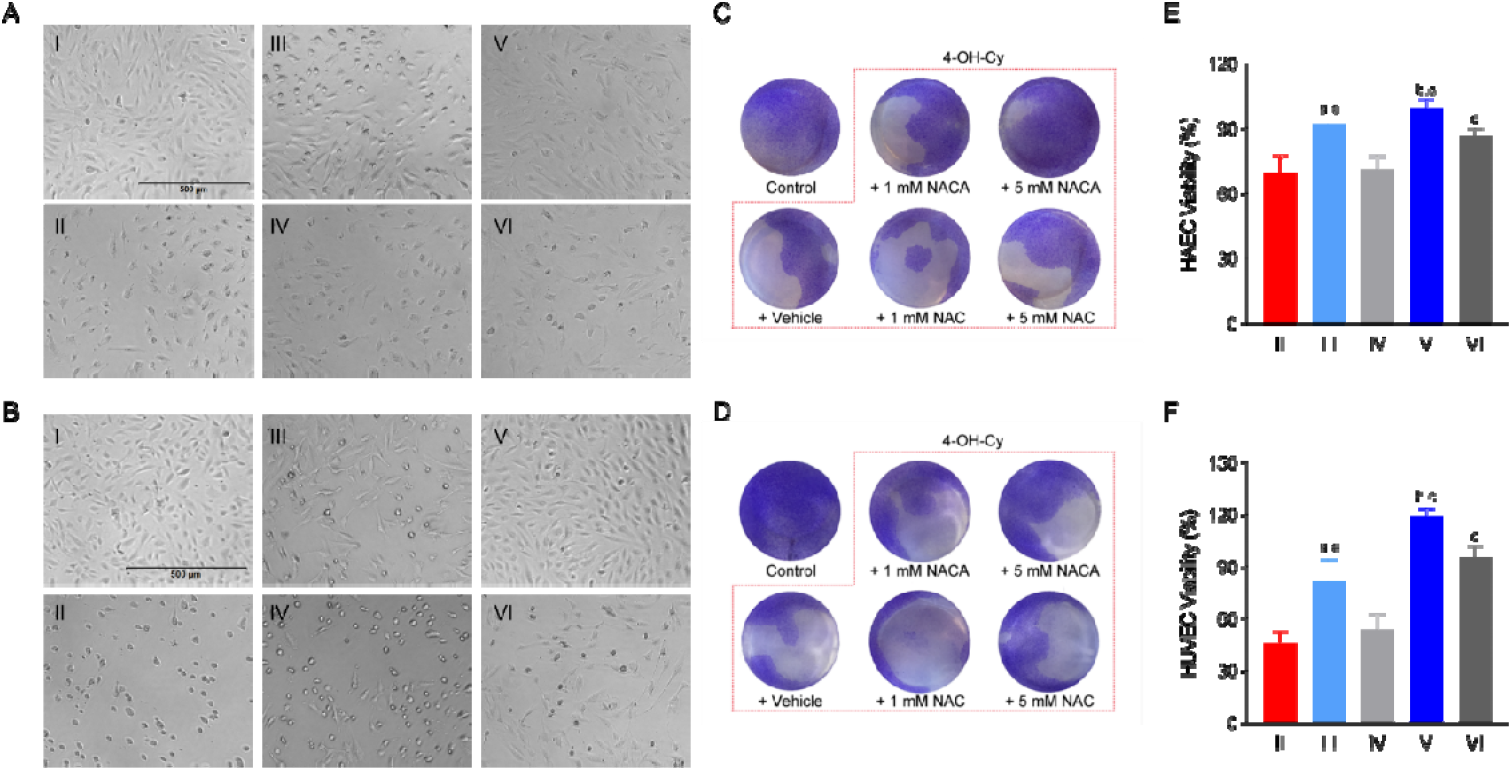
Endothelial cell morphology and viability. Endothelial cells were pre-treated with NACA or NAC for 6 hr. After being rinsed once with HBSS, HAECs were further treated with 32 μM 4-OH-Cy for 20 hr, while HUVECs were incubated with 35 μM 4-OH-Cy for 20 hr. (A-B) The cell morphology of HAEC (A) and HUVEC (B) was imaged by inverted phase contrast microscopy. The scale bar applies to all pictures. I: control; II: 4-OH-Cy; III: 1 mM NACA + 4-OH-Cy; IV: 1 mM NAC + 4-OH-Cy; V: 5 mM NACA + 4-OH-Cy; VI: 5 Mm NAC + 4-OH-Cy. (C-D) HAECs (C) and HUVECs (D) were fixed and subjected to crystal violet staining. (E-F) HAEC (E) and HUVEC (F) cell viability was determined using WST-1 kit. N = 6. The control group represented for cells without treatment and its viability was considered as 100%. a: p < 0.05 when it was compared to IV; b: p < 0.05 when it was compared to VI; c: p < 0.05 when it was compared to II.

### N-acetylcysteine amide attenuates 4-hydroxycyclophosphamide-incuced endothelial cell apoptosis and DNA damage

Basically, apoptosis has been recognized as a pivotal mode of programmed cell death[29], and Cy acts through the formation of irreversible cross-linkage in DNA, leading to cell apoptosis. To detect the efficacy of NACA in reducing apoptosis, HAECs and HUVECs were stained with Annexin V which recognizes the apoptosis marker phosphatidylserine and 7-Amino-Actinomycin (7-AAD) which has a high affinity for DNA. In line with previous studies, we found strong signs of apoptosis in a time and concentration-dependent manner on both types of endothelial cells (Figure S2). To investigate the protective effect of NACA and NAC, endothelial cells were successively incubated with the antioxidants and 4-OH-Cy, and the apoptosis rate was evaluated by the sum percentage of early (Annexin V^+^7-ADD^-^) and late apoptotic (Annexin V^+^7-ADD^+^) cells. As shown in Figure 3A-D, 1 mM of NACA decreased the relative apoptosis rate to 60% for HAECs and 80% for HUVECs, respectively, whereas there was no effect for 1 mM of NAC. When the concentration was raised to 5 mM, both antioxidants demonstrated significant protective capability on endothelial cells, yet NACA was still superior to NAC. Specifically, NAC reduced the relative apoptosis rate to around 60%, while the decrement by NACA was approximately 70%.

**Figure 3.**
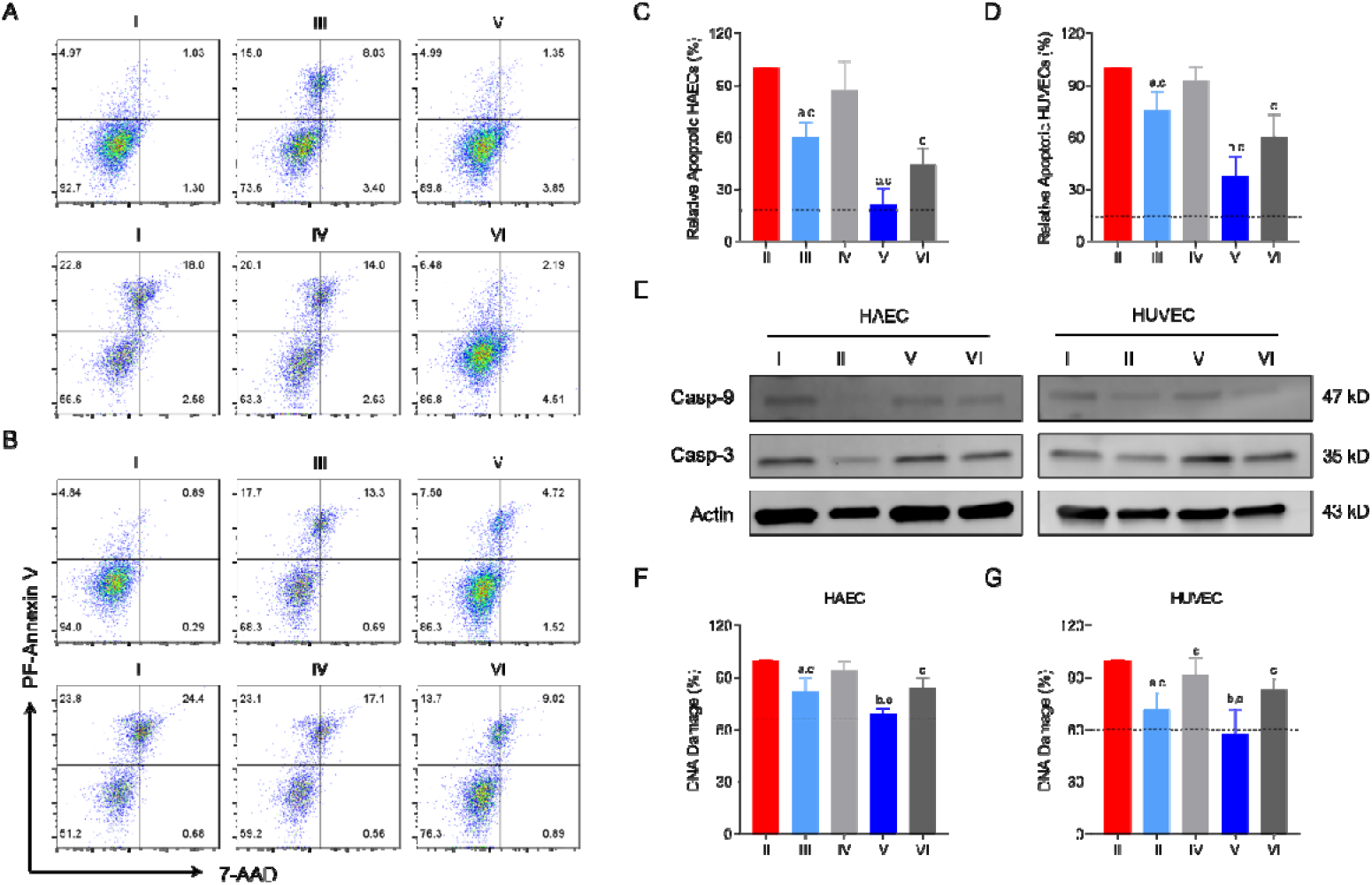
NACA attenuates 4-OH-Cy-induced endothelial cell apoptosis and DNA damage. Endothelial cells were pre-treated with NACA or NAC for 6 hr. After rinsing, HAECs were further treated with 32 μM 4-OH-Cy for 20 hr, while HUVECs were incubated with 35 μM 4-OH-Cy for 20 hr. (A-B) HAECs (A) and HUVECs (B) were stained by PE-Annexin V apoptosis detection kit I. Representative scatter plots from flow cytometry were shown. I: control; II: 4-OH-Cy; III: 1 mM NACA + 4-OH-Cy; IV: 1 mM NAC + 4-OH-Cy; V: 5 mM NACA + 4-OH-Cy; VI: 5 mM NAC + 4-OH-Cy. (C-D) Relative apoptotic cells upon each treatment. Apoptosis rate in the group treated with 4-OH-Cy was set to 100%. Blue dash line refers to the apoptosis rate in control group without treatment. N = 5. (E) Protein level of intact Casp-9 and Casp-3 in the whole cell lysate was determined using western blot. Actin was applied as the reference protein. (F-G) Relative DNA damage in endothelial cells. After treatment, cells were fixed, permeabilized, and stained with recombinant anti-gamma H2A.X (phospho S139) antibody. Data was collected by flow cytometry. Percentage of DNA damage in the group treated with 4-OH-Cy was considered as 100%. Blue dash line refers to the relative DNA damge in control group without treatment. N = 5. a: p < 0.05 when it was compared to IV; b: p < 0.05 when it was compared to VI; c: p < 0.05 when it was compared to II.

Caspase 9 (Casp-9) and caspase 3 (Casp-3) are well-known executors for apoptosis. Cleavage of Casp-9 initiates apoptosis by activating the downstream executioner caspases[30], while activation of Casp-3 is responsible for chromatin condensation and DNA fragmentation[31]. Therefore, the levels of these caspases were further measured in whole cell lysate. After 20 hr treatment of 4-OH-Cy, Casp-9 and Casp-3 was downregulated in HAECs and HUVECs. Remarkably, pre-incubation with 5 mM of NACA or NAC reversed the changes of caspases in both endothelial cell lines, and significant difference between NACA and NAC was observed (Figure 3E, Figure S3).

Apart from cell apoptosis, we also examined the effect of the antioxidants on 4-OH-Cy-induced DNA damage by identification of H2A.X phosphorylation. As depicted in Figure 3F and 3G, NACA and NAC reduced the DNA damage in a dose-dependent manner, and NACA showed a better capability than NAC. It is noteworthy that the DNA damage caused by 4-OH-Cy was completely abrogated by 5 mM of NACA in both HAECs and HUVECs, with an equal percentage as the control group was detected.

### N-acetylcysteine amide confers protection against 4-OH-Cy-induced oxidative stress

Given that NACA is a potent antioxidant, and it showed the ability to prevent endothelial cell from 4-OH-Cy-induced death, DCF-DA probe was exploited to investigate the effect of NACA on cellular ROS alteration following 4-OH-Cy treatment. We found excessive ROS accumulated in HAECs and HUVECs after 4-OH-Cy treatment. In line with our previous results, 1 mM of NAC could not cope with the excessive ROS in 4-OH-Cy-treated cells, while the same concentration of NACA significantly ameliorated ROS accumulation (Figure 4A and 4B).

**Figure 4.**
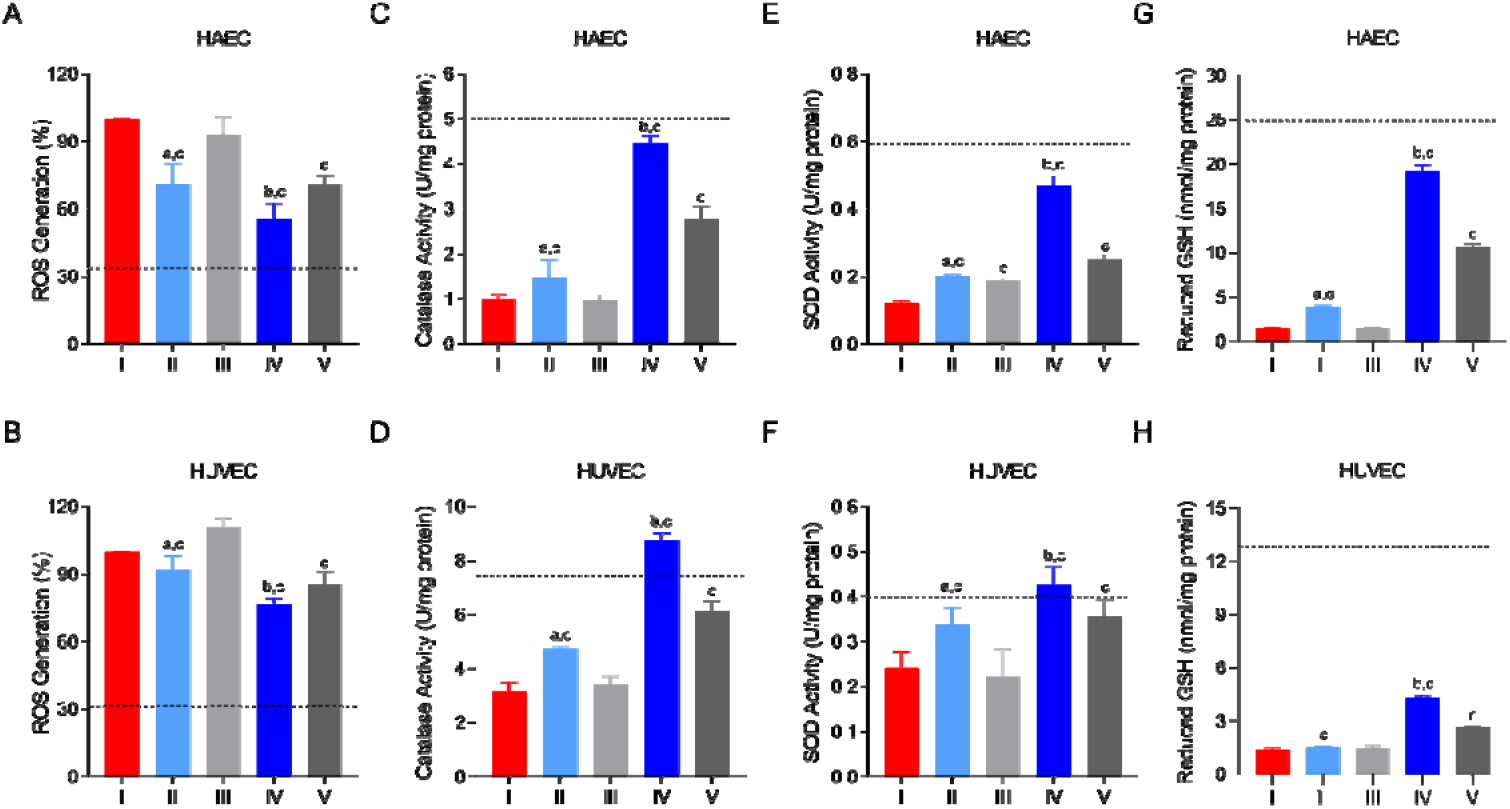
NACA confers protection against 4-OH-Cy-induced oxidative stress. (A-B) ROS levels in HAECs (A) and HUVECs (B) after 4-OH-Cy exposure with or without antioxidants pre-incubation. The amount of ROS generation in the group treated with 4-OH-Cy was set to 100%. N = 5. (C-D) Absolute catalase activity in endothelial cells with different treatments. N = 5. (E-F) Absolute SOD activity in endothelial cells. N = 5. (G-H) The content of intracellular reduced GSH. N = 4. I: 4-OH-Cy; II: 1 mM NACA + 4-OH-Cy; III: 1 mM NAC + 4-OH-Cy; IV: 5 mM NACA + 4-OH-Cy; V: 5 mM NAC + 4-OH-Cy. Blue dash line refers to the level in control group without treatment. a: p < 0.05 when it was compared to III; b: p < 0.05 when it was compared to V; c: p < 0.05 when it was compared to I.

We also assessed the ability of NACA to strengthen the intracellular antioxidative defense system in both HAECs and HUVECs. The activities of two enzymatic antioxidants (i.e., catalase and SOD) and the level of reduced GSH were detected. With exposure to 4-OH-Cy, the activities of both enzymes were compromised to a large extent, especially in HAECs. One mM of NACA significantly enhanced the cellular level of catalase and SOD, whereas NAC was able to show the efficacy at a higher concentration. Strikingly, 5 mM of NACA nearly reinforced catalase and SOD to a normal level, where a 4-fold increment in HAECs and a 2-fold change in HUVECs were noticed compared to the cells only incubated with 4-OH-Cy (Figure 4C-F). The concentration of reduced GSH was examined under different conditions to clarify its contribution. Compared to the cells without treatment, a conspicuous decrease in cellular GSH level was observed in both HAECs (from 24.9 nmol/mg protein to 1.5 nmol/mg protein) and HUVECs (from 6.5 nmol/mg protein to 0.7 nmol/mg protein) following 4-OH-Cy treatment. As GSH precursors, NACA and NAC at a concentration of 5 mM partially replenished the GSH content and NACA exhibited superior ability to NAC on HAECs and HUVECs. These results were in compliance with our previous *in vivo* study[24].

### N-acetylcysteine amide abrogated eNOS/arginase imbalance in endothelial cells

Since both eNOS and arginase imbalance contributes to vascular dysfunction, we evaluated the effects of NACA on the protein expression level of eNOS and arginase in endothelial cells exposed to 4-OH-Cy. Our results revealed that 4-OH-Cy significantly reduced eNOS protein expression and increased the level of arginase I in HAECs compared to the control group, but it increased the expression levels of both proteins in HUVECs. These effects were reversed in the cells treated with 5 mM of NACA, while the effect of NAC were not obvious at the same concentration. Interestingly, neither NACA nor NAC improved the protein expression level of arginase II which was downregulated by 4-OH-Cy (Figure 5).

**Figure 5.**
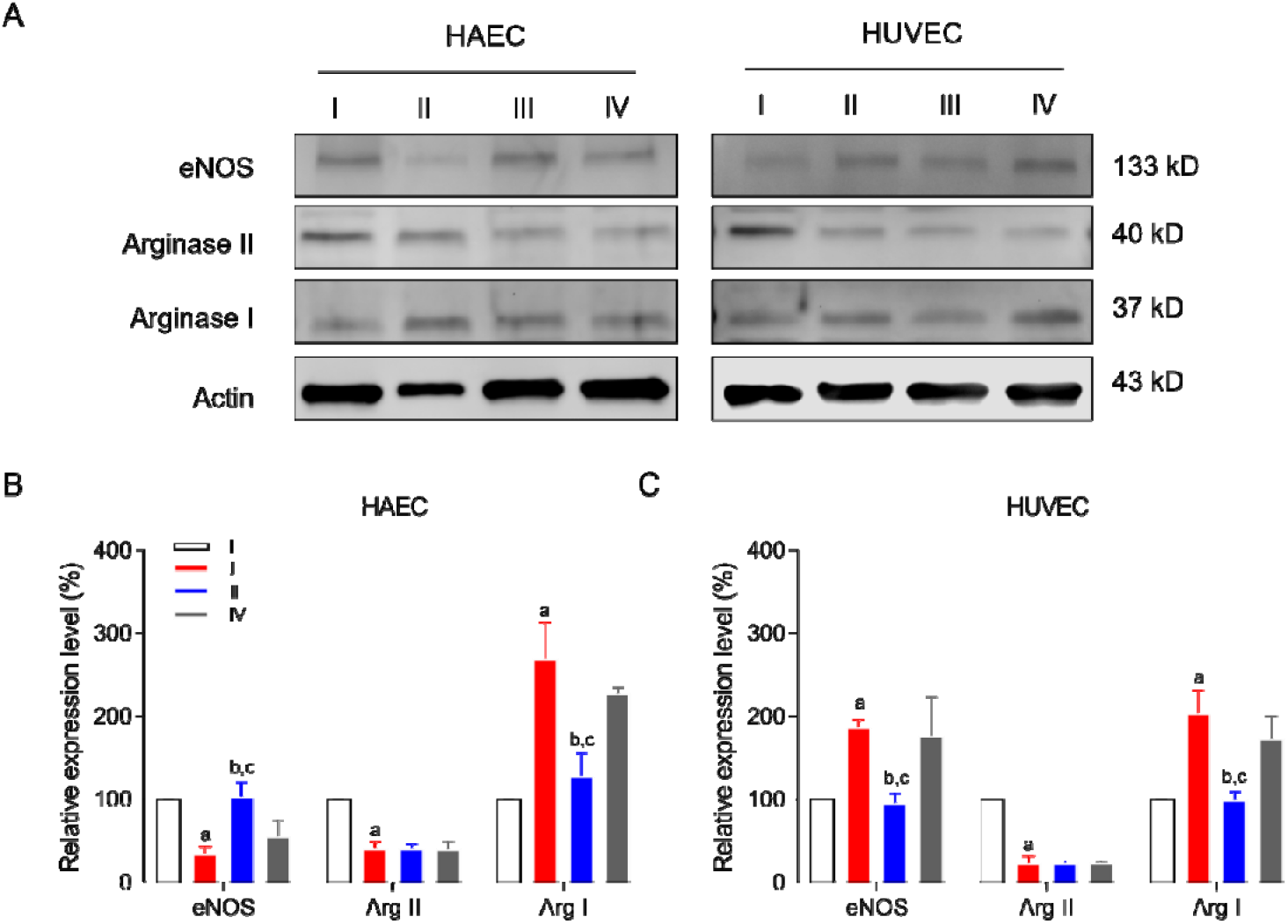
(A) Western blot analysis revealed the expression of eNOS and arginases. (B-C) Relative protein level of eNOS, Arg II, and Arg I in endothelial cells. Endothelial cells were pre-treated with 5 mM of NACA or NAC for 6 hr, then HAECs were further treated with 32 μM 4-OH-Cy for 20 hr, and HUVECs were incubated with 35 μM 4-OH-Cy for 20 hr. Cells were lysed and protein level was measured by western blot. Actin was applied as the reference protein, and the relative expression level is defined as the percentage compared to actin. I: control; II: 4-OH-Cy; III: 5 mM NACA + 4-OH-Cy; IV: 5 mM NAC + 4-OH-Cy. Results are presented as mean ± SD of three independent experiments. a: p < 0.05 when it was compared to I; b: p < 0.05 when it was compared to II. c: p < 0.05 when it was compared to IV.

### N-acetylcysteine amide preserves angiogenesis of endothelial cells

Neovascularization is an essential feature of endothelial cells; therefore, a tube formation assay was performed as an *in vitro* assessment of angiogenic properties. HAECs and HUVECs capillary tubular structure formed on BME matrix was significantly inhibited by 4-OH-Cy, resulting in a shorter total tube length, a less amount of master junctions and segments, a smaller area of meshes, a longer length of isolated branches, and a greater number of extremities (Figure 6A-F; Figure S4). Nevertheless, the antioxidants, especially NACA, restored the angiogenic ability in HAECs and HUVECs. In contrast to the cells only exposed to 4-OH-Cy, those pre-incubated with 1 mM of NACA already demonstrated an increase by 30% in the total tube length and an elevation by 40% in total meshes area. Unsurprisingly, 5 mM of NACA improved the tube formation capability to a level that was comparable with it in the control group. Unlike NACA, 1 mM of NAC only showed a minor enhancement in both endothelial cells, and its effect was enlarged and became significant when the concentration was raised to 5 mM.

**Figure 6.**
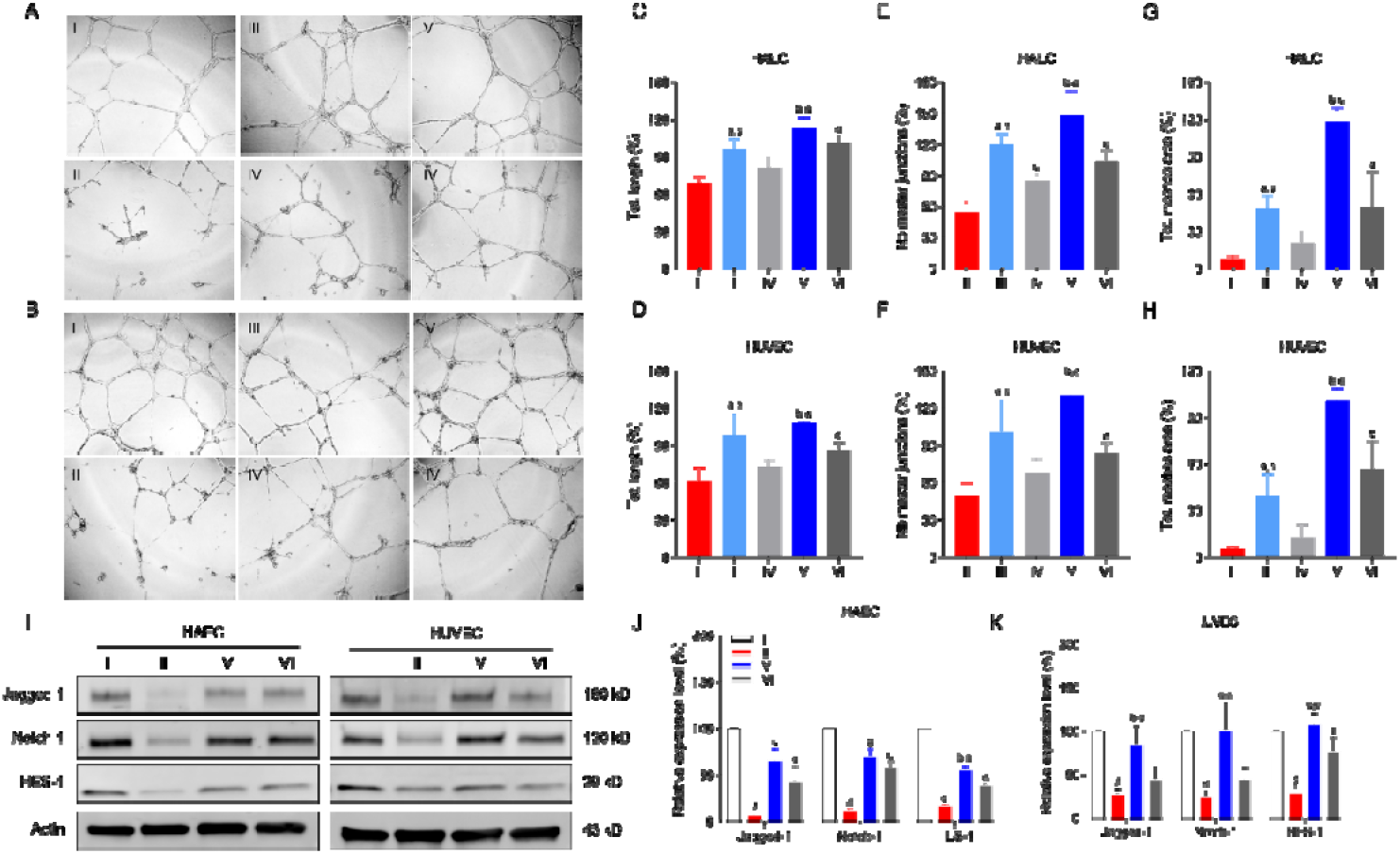
In vitro angiogenesis assay and expression level of Notch-1 Jagged-1 and HES-1 in endothelial cells. (A-B) Representative phase-contrast micrographs of tubular structures in cultured HAECs (A) and HUVECs (B) after different treatments. (C-H) Quantification of total tube length (C&D), number of master junctions (E&F) and total meshes area (G&H) in percent. The data from group I was defined as 100%. N = 5. (I) Western blot analysis for the expression of Jagged-1, Notch-1, and HES-1. (J-K) Relative protein level of Jagged-1, Notch-1, and HES-1 in endothelial cells. N = 3. I: control; II: 4-OH-Cy; III: 1 mM NACA + 4-OH-Cy; IV: 1 mM NAC + 4-OH-Cy; V: 5 mM NACA + 4-OH-Cy; VI: 5 mM NAC + 4-OH-Cy. a: p < 0.05 when it was compared to IV; b: p < 0.05 when it was compared to VI; c: p < 0.05 when it was compared to II; d: p < 0.05 when it was compared to I.

To further explore the underlying protective mechanism of NACA against 4-OH-Cy-induced angiogenesis dysfunction, the Notch signaling was investigated in the present study as it has been proven to be a pivotal factor for sprouting angiogenesis and vessel maturation. As shown in Figure 6G-I, the expression levels of Notch-1, Jagged-1 and their downstream product HES-1 in HAECs and HUVECs were significantly decreased following the exposure to 4-OH-Cy, and this effect was attenuated by pre-treatment with 5 mM of NACA or NAC, yet the difference between NACA and NAC was significant.

## Discussion

Cy is a widely used anti-cancer drug and routinely involved in the regimens of HCT. Although Cy is relatively well tolerated at lower doses, high-dose regimens such as those given prior to or post HCT can be associated with a variety of cardiacvascular adverse events[32]. Most studies exploring the Cy-induced cardiotoxicity have focused on cardiomyocytes. Endothelial cells constitute the inner layer of blood vessels and cardiac chamber and bear the ability for thromboresistance, regulation of cellular adhesion and maintenance of vascular contraction and dilation[33], however, relevant studies on endothelium in context of Cy-induced cardiotoxicity are lacking. Our current work has contributed a comprehensive and designated insight into the endothelial damage originated from Cy exposure. In agreement with previous reports[34, 35], we confirmed the presence of ICAM-1 upregulation, oxidative stress and eNOS uncoupling in Cy-induced endothelial injury. Moreover, we demonstrated that blockage of Notch signaling pathway is another underlying mechanism, and compared to the parent compound NAC, the novel antioxidant NACA displayed higher efficacy against Cy-induced endothelial dysfunction.

In our mouse model, significant circulating cTNI elevation was observed after a single high dose of Cy. In line with our finding, a clinical study also revealed that patients with suspected coronary endothelial dysfunction were associated with higher cTNI levels[26]. ICAM-1 is constitutively present on endothelial cells and is one of the major adhesion molecules mediating leukocytes recruitment from circulation to inflammatory sites[36]. The expression of ICAM-1 is upregulated by proinflammatory cytokines, and its increased secretion causes local endothelial cell leakiness and activation[37]. Cy has been reported to induce multi-organ toxicity by generating excessive ROS via NF-κB/TNF-α pathway[38], and the same pathway was coincidentally shared by the ICAM-1 overexpression[39, 40]. Therefore, we hypothesized that NACA, as a ROS scavenger, might hinder upregulation of ICAM-1 and subsequent endothelial injury. Indeed, a clear enhancement of local ICAM-1 expression was observed after Cy treatment, and NACA was shown to be effective in both downregulation ICAM-1 and maintain the aortic endothelium morphologically.

Results from our *in vitro* studies showed that 4-OH-Cy treatment significantly induced endothelial cell death by stimulating apoptosis, with the signs of outer membrane blebbing, positive Annexin/7-ADD staining, diminished Casp-3/9, and DNA damage (Figure 2A-B; Figure 3; Figure S2), which is consistent with other studies[41, 42]. 4-OH-Cy, the active metabolite of Cy, is further metabolized to aldophosphamide and results in the formation of phosphoramide mustard and acrolein via β elimination[43]. Besides the direct effect of phosphoramide mustard on DNA, acrolein could drive ROS generation and indirectly causes DNA damage, lipid peroxidation and cell apoptosis[44]. Therefore, the benefits from NACA/NAC might originate from their effects on the acrolein pathway and partially, but not completely, revere Cy-derived toxicity.

Cells possess an antioxidative system, including enzymes and non-enzymatic substances, to cope with the detrimental effect of excessive ROS like superoxide anion (O_2_ ^-^), hydrogen peroxide (H_2_O_2_), and hydroxyl radical (HO•)[45]. Oxidative stress occurs when the endogenous antioxidant defense system is overwhelmed by ROS through either a decreased ability of cellular antioxidation or an increased level of ROS. In line with previous studies[7, 35], the oxidative stress induced in endothelial cells with Cy was evident by dramatically elevated intracellular ROS level, decreased activities of SOD and catalase, and declined content of GSH (Figure 4). SOD catalyzes O_2_^-^ into oxygen and H_2_O_2_, while catalase acts as a scavenger to convert H_2_O_2_ into oxygen and H_2_O. On the other hand, GSH is the most abundant reducing equivalent that neutralizes ROS[46] and in the meanwhile covalently modifies cytostatic compounds like 4-OH-Cy for detoxification[47]. As a ROS scavenger and a GSH precursor, either NAC or NACA showed the capability to combat with the oxidative stress in endothelial cells. Nevertheless, due to better cell permeability and ability to replenish low molecular mass thiols (e.g., cysteine, homocysteine, cysteinylglycine and GSH)[24, 48-50], NACA exhibited higher efficacy in defending oxidative stress compared to NAC (Figure 4).

Vascular tone is critical for endothelial hemostasis and largely dependent on the availability of NO. NO synthesis is accomplished by eNOS along with its co-factor tetrahydrobiopterin (BH_4_) using L-arginine as the substrate, however, L-arginine is competed by arginase as well to form ornithine and urea. Deficiency in either substrate or co-factor contributes to uncoupled eNOS and generation of superoxide and peroxynitrite (ONOO^-^) rather than NO. Moreover, ROS can exhaust L-arginine via upregulating arginase activity and aggravates endothelial dysfunction[51]. In the present study, we found that the expression level of arginase I was notably upregulated in both endothelial cells upon 4-OH-Cy treatment, implying the escalated consumption of L-arginine. Noteworthy, NACA pretreatment significantly reversed the effect from 4-OH-Cy while NAC did not. In contrast, 4-OH-Cy impeded arginase II level that has not been influenced by NACA or NAC. The underlying mechanism is unclear but, it might be due to compensation of increased arginase I. Intriguingly, our results showed different trends of eNOS alteration in endothelial cells, where a downregulation in HAECs and an upregulation in HUVECs were seen. Previous studies have implied that decreased eNOS contributes to NO scarcity whereas increased eNOS leads to excessive NO and nitrative stress[52, 53]. Regardless of the alteration patterns, NACA normalized eNOS to a basal level and it effect was superior to NAC.

Formation of new vessels is another fundamental trait of endothelium[54]. Using a tube formation assay, we validated that 4-OH-Cy suppressed the angiogenetic behavior of endothelial cells *in vitro*, and the suppression was abrogated by NACA and NAC to different extents (Figure 6). In search for a plausible explanation, Notch signaling appeared in our sight for its crucial role in modulating angiogenesis[55]. In principle, the Notch signaling pathway is initiated when Notch ligand (e.g., Jagged-1) binds to Notch receptor (e.g., Notch-1), and consequent outcome is induction of the transcriptional process of target genes (e.g., Hes-1) [56]. On the basis of this notion, we found that exposure of 4-OH-Cy significantly stifled the Notch1 pathway activation in HAECs and HUVECs (Figure 6). It has been agreed that intracellular inhibition of Notch signaling and increased ROS level are mutual cause and effect [54]. By pretreating the endothelial cells with NACA or NAC, we validated that expression level of proteins involved in Notch1 signaling has been significantly preserved through scavenging ROS (Figure 6).

## Conclusion

This study proved that the novel antioxidant NACA could be applied to preventing Cy-induced endothelial dysfunction. The preventive effect was mediated through reduction of oxidative stress, inhibiting cell apoptosis, maintenance of eNOS/arginase balance, and orchestrating ICAM-1 expression. In the meanwhile, NACA recovered the angiogenic ability of endothelial cells via regulating Notch-1 signaling pathway. In comparison to NAC, NACA appears to be a better prophylactic agent for Cy-induced endothelial toxicity, suggesting lower doses and shorter treatment duration may be needed to achieve similar efficacy. Our findings will instruct further investigation to apply NACA as a potential prophylactic and therapeutical medication.

## Acknowledgement

This study was supported by grants from the Swedish Research Council (2017-00741), Cancer Society (CAN2014/759), Barncancer fonden (PR2017-0083), KI funds (2018-02377), and Cancer Research Funds of Radiumhemmet project (161082) to M.H.. R.H. receives the PhD student scholarship from China Scholarship Council.

## Contributions

R.H., Y.Z., and M.H. conceived the study. R.H., W.Z., T.S., and Y.Z performed experiments and analyzed data. M.H. supervised the study and acquired funding. R.H., W.Z., Y.Z. and M.H. wrote the manuscript. All authors participated in the preparation of the manuscript.

## Supporting Information

Figure S1. Cytotoxicity of 4-hydroxy-cyclophosphamide, N-acetylcysteine, and N-acetylcysteine amide on endothelial cells.

Figure S2. Endothelial cell apoptosis induced by 4-hydroxy-cyclophosphamide.

Figure S3. Protein level of Caspase-3 and Caspase-9 in endothelial cells.

Figure S4. Angiogenic parameters obtained from tube formation assay.

