## supplementary materials for "A novel antioxidant N-acetylcysteine Amide Alleviates Cyclophosphamide-induced Endothelial Damage"

**Figure S1. Cytotoxicity of 4-hydroxy-cyclophosphamide, N-acetylcysteine, and N-acetylcysteine amide on endothelial cells.** (A) HAECs and HUVECs were treated with different concentrations of 4-OH-Cy for 20 hr and cell viability was determined using WST-1 kit. (B) Cytotoxicity of NAC on HAECs and HUVECs after 6 hr treatment. Cell viability was determined by Cell Titer-Glo kit (Promega, G9241). (C) Effect of NACA on HAECs and HUVECs after 6 hr treatment. Cell viability was determined by CellTiter-Glo kit. N = 3.

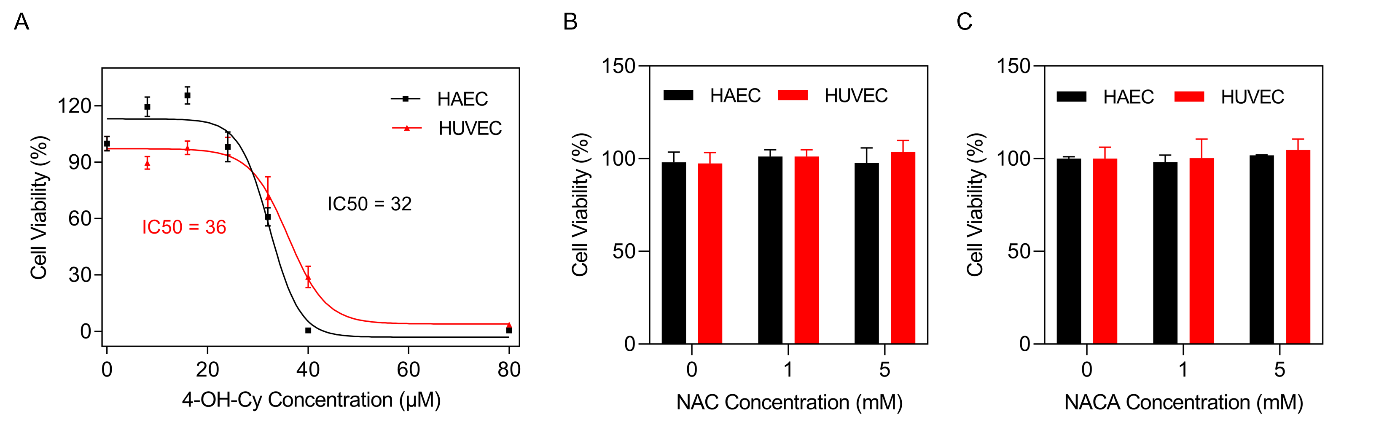

**Figure S2. Endothelial cell apoptosis induced by 4-hydroxy-cyclophosphamide.** (A) HAECs were treated with 60 µM of 4-OH-Cy for 6 hr, 8.5 hr and 11 hr, and then stained using PE-Annexin V apoptosis detection kit I. Representative scatter plots from flow cytometry were shown. (B) HUVECs were treated with 80 µM of 4-OH-Cy for 1 hr, 3 hr and 5 hr, and then stained for apoptosis analysis. Representative scatter plots from flow cytometry were shown. (C-D) HAECs (C) and HUVECs (D) were treated with 15 µM, 35 µM and 50µM of 4-OH-Cy for 20 hr, and then stained for apoptosis analysis. (E-H) Quantification of cell statues corresponding to A-D, respectively.

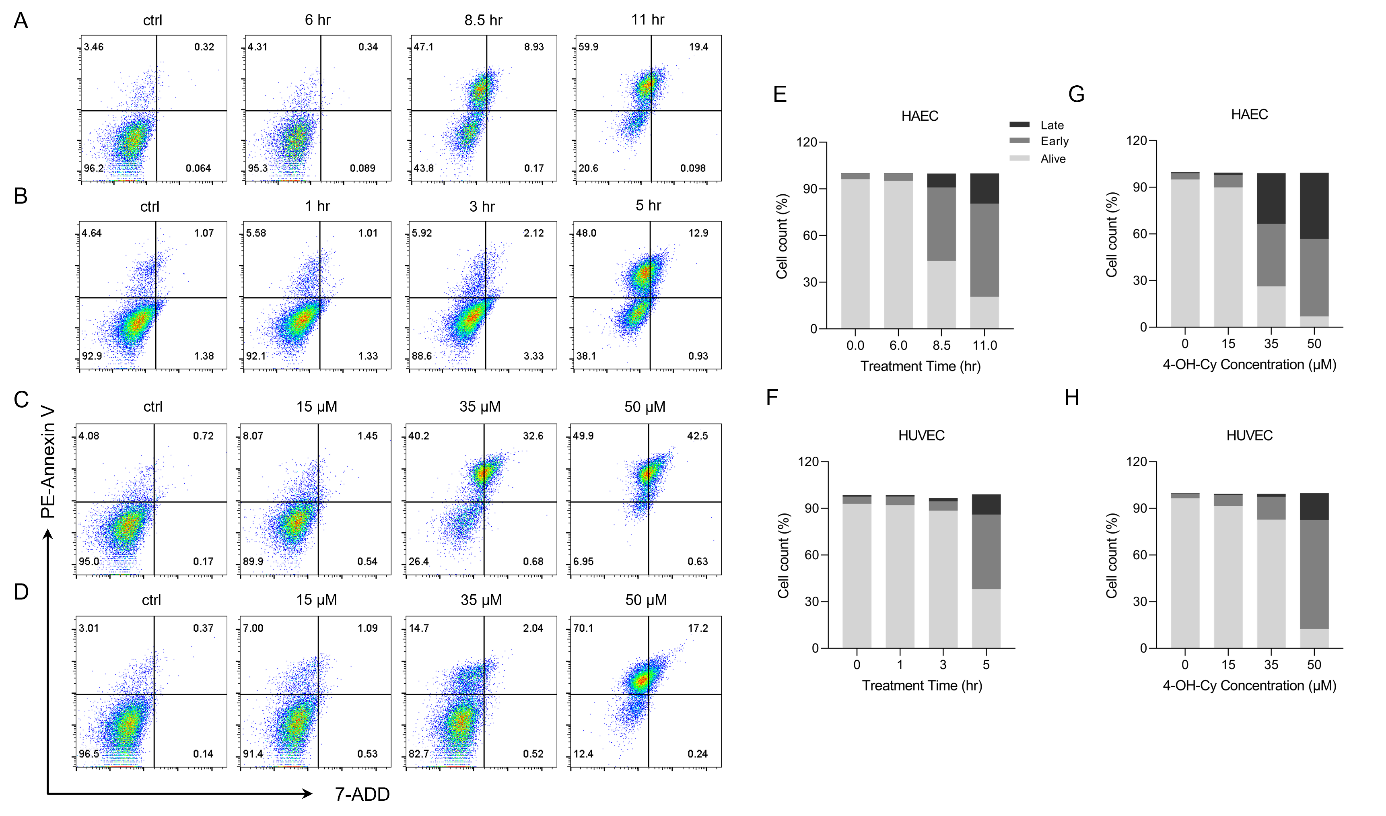

**Figure S3. Protein level of Caspase-3 and Caspase-9 in endothelial cells.** Endothelial cells were pre-treated with 5 mM of NACA or NAC for 6 hr. After being rinsed once with HBSS, HAECs (A) were further treated with 32 µM 4-OH-Cy for 20 hr, while HUVECs (B) were incubated with 35 µM 4-OH-Cy for 20 hr. Cells were lysed and the protein level was measured by western blot. Actin was applied as the reference protein, and the relative expression level is defined as the percentage compared to actin. I: control; II: 4-OH-Cy; III: 5 mM NACA + 4-OH-Cy; IV: 5 mM NAC + 4-OH-Cy. Results are presented as mean ± SD of three independent experiments. a: p < 0.05 when it was compared to I; b: p < 0.05 when it was compared to II; c: p < 0.05 when it was compared to VI.

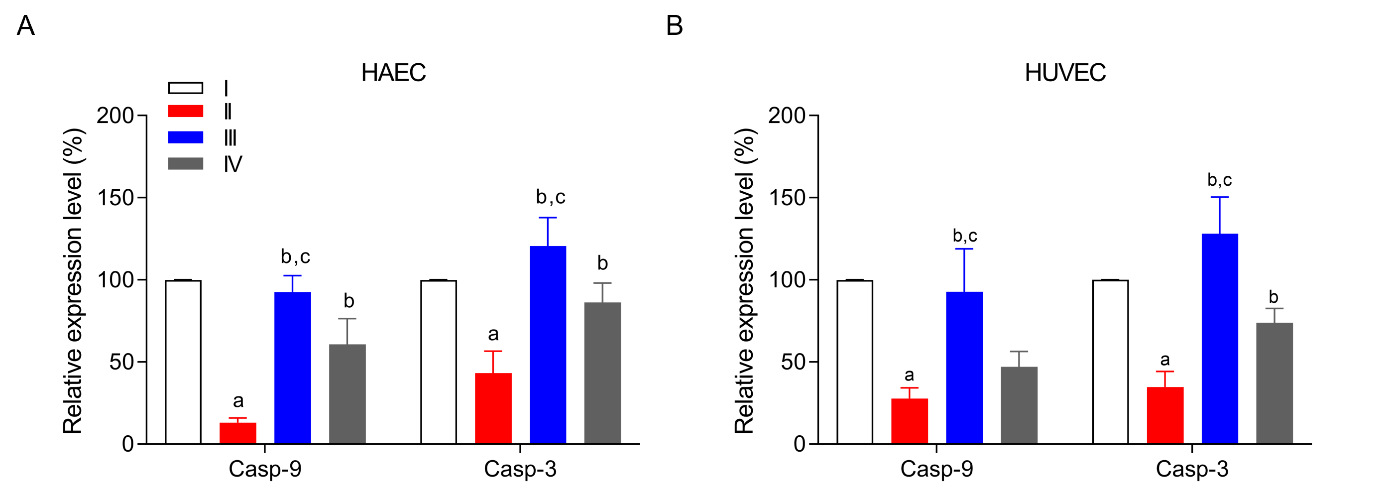

**Figure S4. Angiogenic parameters obtained from tube formation assay.** Quantification of (A-B) relative master segments number, (C-D) relative number of meshed, (E-F) relative total length of isolated branches and (G-H) relative number of extremities in culture HAEC and HUVEC after different treatments. Calculations were performed by Image J software. The data from control group without treatment was defined as 100%. N = 5. I: 4-OH-Cy; II: 1 mM NACA + 4-OH-Cy; III: 1 mM NAC + 4-OH-Cy; IV: 5 mM NACA + 4-OH-Cy; V: 5 mM NAC + 4-OH-Cy. a: p < 0.05 when it was compared to III; b: p < 0.05 when it was compared to V; c: p < 0.05 when it was compared to I.

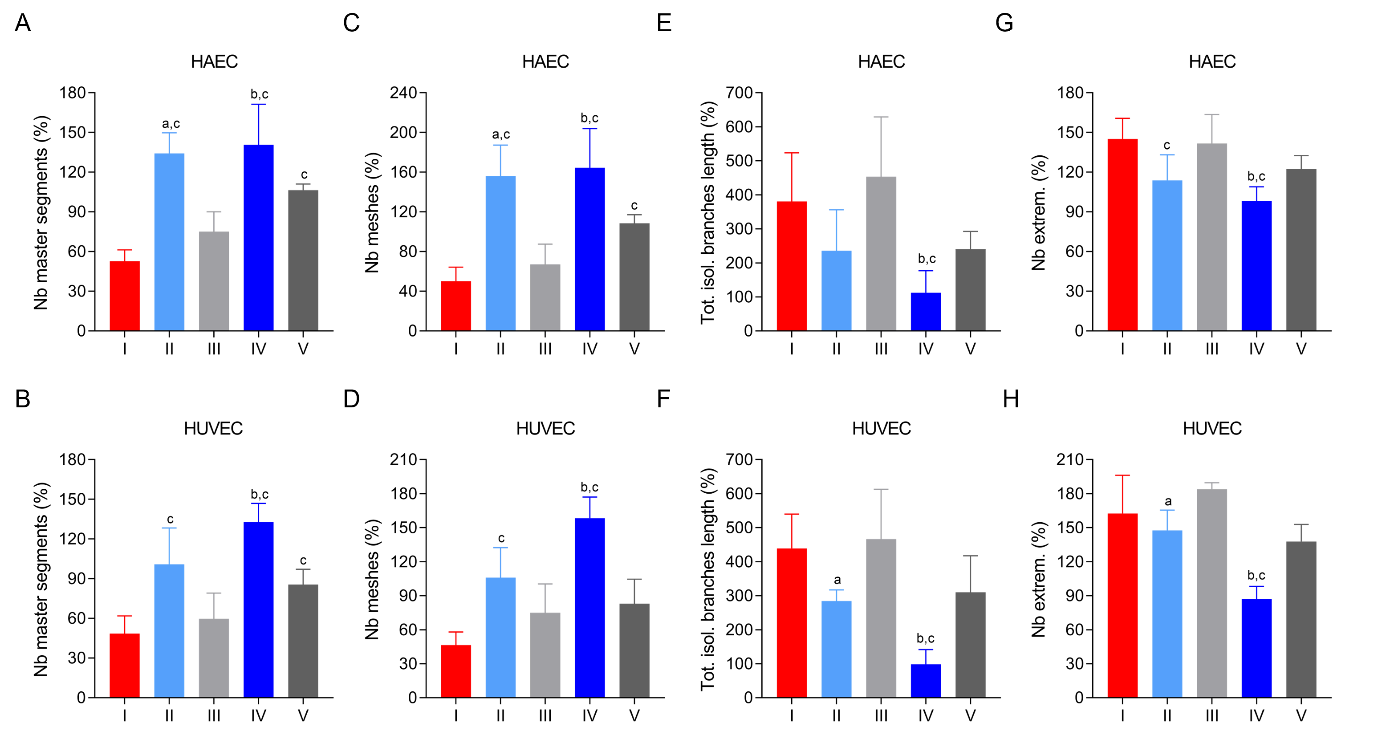
